# GGDB: A Grameneae Genome Alignment Database of Homologous Genes Hierarchically Related to Evolutionary Events

**DOI:** 10.1101/2022.01.20.477137

**Authors:** Qihang Yang, Tao Liu, Tong Wu, Tianyu Lei, Yuxian Li, Xiyin Wang

## Abstract

Owing to their economic values, Gramineae plants have been preferentially sequenced their genomes. These genomes are often quite complex, e.g., harboring many duplicated genes, which were the main source of genetic innovation and often the results of recurrent polyploidization. Deciphering the complex genome structure and linking duplicated genes to specific polyploidization events are important to understand the biology and evolution of plants. However, the effort has been held back due to its high complexity in analyzing these genomes. Here, by hierarchically relating duplicated genes in colinearity to each polyploidization or speciation event, we analyzed 29 well-assembled and up-to-date Gramineae genome sequences, separated duplicated genes produced by each event, established lists of paralogous and orthologous genes, and eventually constructed an on-line database, GGDB (http://www.grassgenome.com/). Homologous gene lists from each plant and between them can be displayed, searched, and downloaded from the database. Interactive comparison tools were deployed to demonstrate homology among user-selected plants, to draw genome-scale or local alignment figures, phylogenetic trees of genes corrected by exploiting gene colinearity, etc. Using these tools and figures, users can easily observe genome structural changes, and explore the effects of paleo-polyploidy on crop genome structure and function. The GGDB will be a useful platform to improve understanding the genome changes and functional innovation of Gramineae plants.

**Key points:**

1. GGDB is the only portal hosting Grameneae colinear homologous genes hierarchically related to evolutionary events, especially polyploidization, which have occurred recursively.
2. Allows systematic analysis of colinear gene relationships and function origination and/or divergence across Grameneae plants.
3. Serving the Grameneae research community, with new genomes, modules, tools, and analysis.

## INTRODUCTION

Gramineae is a large group of monocotyledonous flowering plants, which can be divided into more than 620 genera and more than 10000 species, covering 20% of the land area of the earth^[1]^. Gramineae contains many important food crops, such as wheat (*Triticum aestivum*), rice (*Oryza sativa*), corn (*Zea mays*), sorghum (*Sorghum bicolor*), and so on^[2–5]^. With the tens of Gramineae genomes being sequenced, it provides a solid data basis for in-depth analysis of functional innovation and evolution of Gramineae genomes.

The study of the Gramineae genomes revealed repeated polyploidy events during the evolutionary history ^[6–8]^. Polyploidy is an abrupt event, which can create a new species with doubled number of chromosomes, produce a large number of repetitive genes^[9–11]^, trigger large-scale reorganization of biological functions, such as regulatory network re-programming and debugging. Polyploidy leads to genomic instability, and a considerable amount of gene loss may occur^[12; 13]^. Gramineae plants could be taken as a good example in that their common ancestor was affected by a tetraploidization ~100 million years ago, followed by the fast originization and divergence of derivative plants^[14–16]^. Gramineae crops, such as wheat and maize, were resulted from further polyploidization event(s). Recursive polyploidization and genome reorganization makes their genomes rather complex ^[17–19]^.

Gene colinearity provides precious means to study complex genomic structures. In extant genomes, a considerable number of colinear genes produced by polyploidy have been retained ^[20]^. In rice genome, there still remain thousands of colinear genes, resulted from the Gramineae-common tetraploidization ^[21; 22]^. The analysis of colinear genes helps identify ancient polyploidy events, deduce the scale and time of their occurrence, and infer gene functional changes ^[23; 24]^.

In that the importance of gene colinearity, the information was often inferred and stored in biological databases, such as JCVI, PGDD, and COGE^[25–27]^. However, the gene colinearity information in these existing databases has non-negligible shortcomings. First, colinear genes were not related to specific polyploidy events due to methodological difficulty to analyze these recursive events. Second, only two or three species were involved to infer gene colinearity, hampering the efforts to do genus- or family-scale level evolutionary or functional analysis.

Here, by integrating approaches of homologous gene dotplotting to compare genomes, characterizing divergence levels of colinear gene homologs, and checking complement patterns of chromosome breakages, colinear homologs produced by different polyploidies (or speciation) could be separated, and hierarchically related to each relative event ^[28; 29]^. Then, we implemented the above mentioned hierarchical gene colinearity inference approaches with the Gramineae plants, and thus using the results established the Gramineae Genome Alignment Database GGDB (http://www.grassgenome.com/), which contains gene colinearity information to explore the chromosome changes, genomic repatterning, and the actual phylogeny of duplicated genes^[30; 31]^ at the Gramineae family-scale level.

## MATERIALS AND METHODS

We summarizes the species information contained in GGDB, the way of data processing, and the composition structure of webpages (Fig.1A-B). The database is constructed by using MySQL to store analysis results^[32–34]^, such as colinear gene lists, chromosome homology within a genome or between genomes, figures to show homolog between genomes, etc. The website was developed by using HTML and PHP. Data was analyzed by using scripts developed by Python and R^[35–38]^.

**Figure 1.**
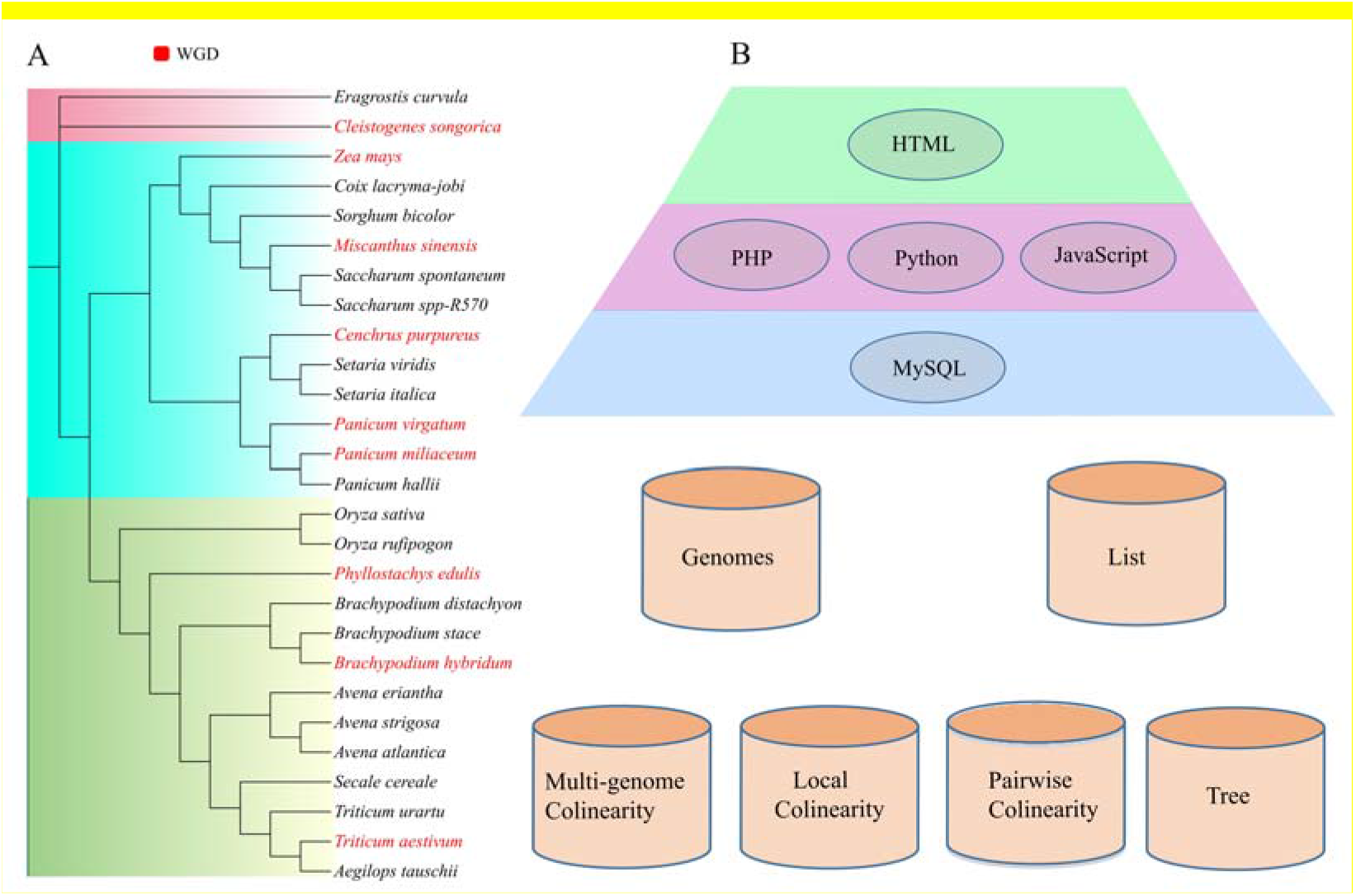
Composition structure of GGDB database. A. A phylogenetic tree of Gramineae species involved in the database; B. the computer languages used to set up tiers of the database and the functional interfaces.

### Data sources

In determining whether a species should be included in the GGDB or not, we require its genome to be assembled to the chromosome-level, allowing credible gene colinearity inference. If multiple genome assemblies are available, we use the latest assembly version in the database^[39–41]^. Most genome data was downloaded from the NCBI database (https://www.ncbi.nlm.nih.gov/) and the JGI database (https://phytozome-next.jgi.doe.gov/) (Table 1). Genome data was preprocessed by using home-made Perl scripts: (i) uniform format of gene names was adopted; (ii) redundant information in the genome annotation is removed; (iii) gene location files was extracted, including the information of chromosome numbers, gene IDs, gene locations and orders on chromosomes, etc.

**Table 1.**
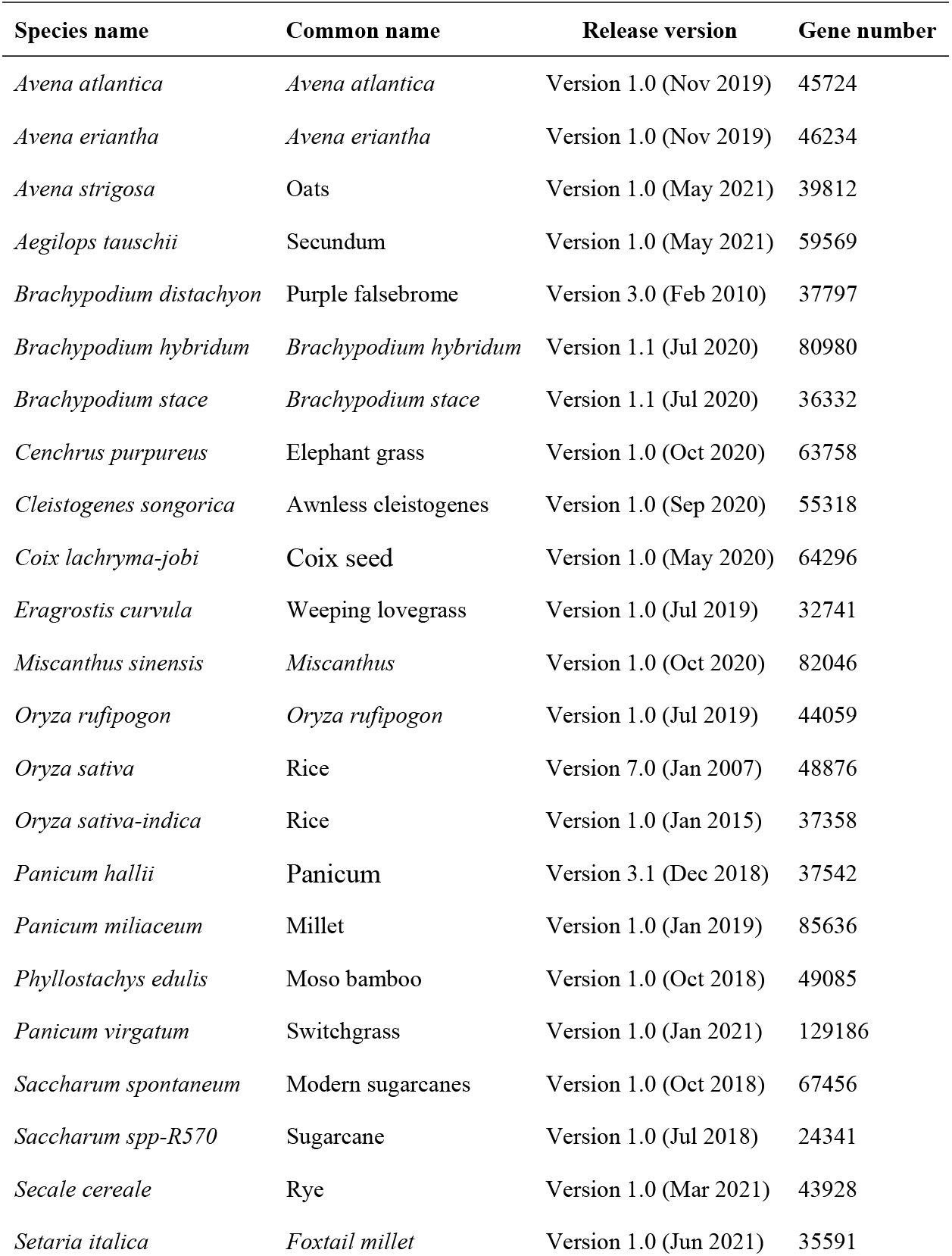
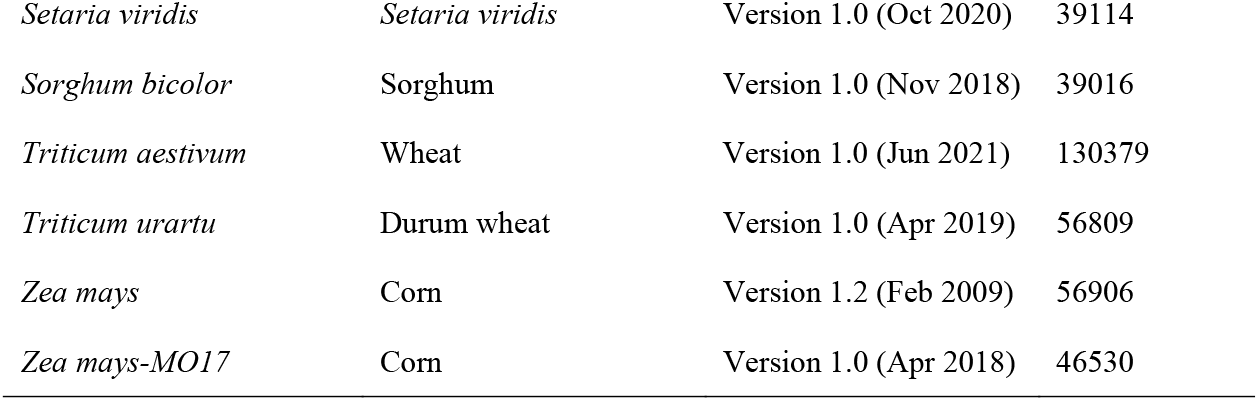
29 plants currently involved in the GGDB.

| Species name | Common name | Release version | Gene number |
| --- | --- | --- | --- |
| *Avena atlantica* | *Avena atlantica* | Version 1.0 (Nov 2019) | 45724 |
| *Avena eriantha* | *Avena eriantha* | Version 1.0 (Nov 2019) | 46234 |
| *Avena strigosa* | Oats | Version 1.0 (May 2021) | 39812 |
| *Aegilops tauschii* | Secundum | Version 1.0 (May 2021) | 59569 |
| *Brachypodium distachyon* | Purple falsebrome | Version 3.0 (Feb 2010) | 37797 |
| *Brachypodium hybridum* | *Brachypodium hybridum* | Version 1.1 (Jul 2020) | 80980 |
| *Brachypodium stace* | *Brachypodium stace* | Version 1.1 (Jul 2020) | 36332 |
| *Cenchrus purpureus* | Elephant grass | Version 1.0 (Oct 2020) | 63758 |
| *Cleistogenes songorica* | Awnless cleistogenes | Version 1.0 (Sep 2020) | 55318 |
| *Coix lachryma-jobi* | Coix seed | Version 1.0 (May 2020) | 64296 |
| *Eragrostis curvula* | Weeping lovegrass | Version 1.0 (Jul 2019) | 32741 |
| *Miscanthus sinensis* | *Miscanthus* | Version 1.0 (Oct 2020) | 82046 |
| *Oryza rufipogon* | *Oryza rufipogon* | Version 1.0 (Jul 2019) | 44059 |
| *Oryza sativa* | Rice | Version 7.0 (Jan 2007) | 48876 |
| *Oryza sativa-indica* | Rice | Version 1.0 (Jan 2015) | 37358 |
| *Panicum hallii* | Panicum | Version 3.1 (Dec 2018) | 37542 |
| *Panicum miliaceum* | Millet | Version 1.0 (Jan 2019) | 85636 |
| *Phyllostachys edulis* | Moso bamboo | Version 1.0 (Oct 2018) | 49085 |
| *Panicum virgatum* | Switchgrass | Version 1.0 (Jan 2021) | 129186 |
| *Saccharum spontaneum* | Modern sugarcane | Version 1.0 (Oct 2018) | 67456 |
| *Saccharum spp-R570* | Sugarcane | Version 1.0 (Jul 2018) | 24341 |
| *Secale cereale* | Rye | Version 1.0 (Mar 2021) | 43928 |
| *Setaria italica* | Foxtail millet | Version 1.0 (Jun 2021) | 35591 |
| *Setaria viridis* | *Setaria viridis* | Version 1.0 (Oct 2020) | 39114 |
| *Sorghum bicolor* | Sorghum | Version 1.0 (Nov 2018) | 39016 |
| *Triticum aestivum* | Wheat | Version 1.0 (Jun 2021) | 130379 |
| *Triticum urartu* | Durum wheat | Version 1.0 (Apr 2019) | 56809 |
| *Zea mays* | Corn | Version 1.2 (Feb 2009) | 56906 |
| *Zea mays-MO17* | Corn | Version 1.0 (Apr 2018) | 46530 |

### Inferring colinear genes

Sequence similarity alignment software BLAST+ was used to infer putative homologous genes (BLAST E-value <= 10^−5^ and sequence matching score >= 100). Homologous gene dotplots were drawn, in that they are helpful to show and infer homology and structural changes within a genome or between genomes. In a homologous gene dotplot, dots represent homologous gene pairs and are often assigned with different colors, to show sequence divergence levels between compared homologous genes.

Colinear genes within and between genomes of Gramineae were inferred by using software ColinearScan^[42]^, using the above BLAST inferred putative homologs, with BLAST accumulated hit scores >= 50 of homologous blocks with colinear gene numbers >= 5, and the statistical significance is set to be <= 1 × 10^−10^ estimated by ColinearScan.

To distinguish colinear gene blocks produced by different events, polyploidies or speciation, we estimated the synonymous nucleic acid substitution rate at the synonymous sites (Ks), which could be used to measure divergence levels between homologous genes. The Nei-Gojobori method implemented in PAML package was used to estimate the above values^[43]^. Actually, homologous blocks produced by different events, especially two polyploidy events, could be mixed together due to having similar Ks values, complicated by the fact that genes evolve at divergent rates. We checked whether homologous blocks shared the chromosome breakage points, which is a hallmark to show them to have been produced by the same polyploidy event. These shared chromosome breakage points could be identified in homologous gene dotplots. When dealing with cross-species gene colienarity, orthologous genes between species often form much better gene collinearity and have much smaller Ks than the outparalogous genes do produced by polyploidy in the common ancestor. Seldom cases need to check chromosome breakage points to distinguish orthologous and outparalogous blocks. Eventually, lists of colinear genes associated with specific polyploidy events or speciation were generated and stored in the MySQL database.

### Multi-genome alignment map

Multi-genome alignment maps were constructed by integrating lists of colinear genes between any two species. A reference genome was selected, and then another plant genomes were aligned to it one by one, with colinear genes as markers. Eventually, a table containing aligned colinear genes was produced, and a gene in the reference genome often has no corresponding homolog in another genome or in the duplicated regions, the corresponding cell in the table were filled with dots.

### Local colinear alignment

Local colinear alignment was constructed by integrating the list of colinear genes among species. We used a Python script to call the integration package Matplotlib module, built a two-dimensional atlas to show gene homology information. Based on the genome comparison software MCScanX[44], we developed the “Local colinear alignment” module. Compared to previous databases, by checking chromosome breakage points, GGDB can distinguish subgenomes produced by genome doubling. After receiving the query data and parameters sent from the web interface, the GGDB server queries the colinear gene results in the database and draws the local colinear alignment figure. The final result file is to be packaged and sent to the browser in PDF format.

### Gene evolutionary tree

To help explore the evolution of duplicated genes, which is critical in genetic innovation, we used phylogenetic analysis software IQTREE, MUSCLE, and FASTTREE^[45–47]^, to construct an evolutionary tree using DNA or protein sequences of a set of homologous genes. Actually, previous research found that duplicated genes, especially those produced by polyploidies, could form trees inconsistent to their real evolutionary relationship^[48]^. For a set of homologous genes, we can construct the expected tree reflecting the actual relationship of colinear genes, including paralogs produced by specific polyploidies, and orthologs originated from speciation.

After accepting the user query parameters (Selected species, reference species, Gene ID) from the browser, the GGDB server queries the database for the homologous colinear genes and writes the gene sequences into a fasta file. The file is used as input for the software MUSCLE to do sequence alignment, and then a tree is built by using FASTTREE^[49; 50]^. Default parameters of these software were used. Both the sequence-alignment derived tree and the expected tree are stored in nwk format, convenient for the user download and further editing. The software EChart plug-in is implemented in JavaScript in our interface, and the interaction between the user and the server is realized^[51; 52]^.

### Construction and content

We developed the GGDB database to provide homologous colinear gene information within each of or between Gramineae plants. The database is currently installed on the CentOS operating system. It has a three-tier architecture, namely, the client tier, the middle tier and the database tier. The client layer that users directly access is developed using PHP and JavaScript. In the database layer, GGDB-related data is stored in a MySQL database. The middle tier receives HTTP requests and is processed by a Apacheweb server. In addition, we include different levels of genomic colinearity analysis tools (Fig.2).

**Figure 2.**
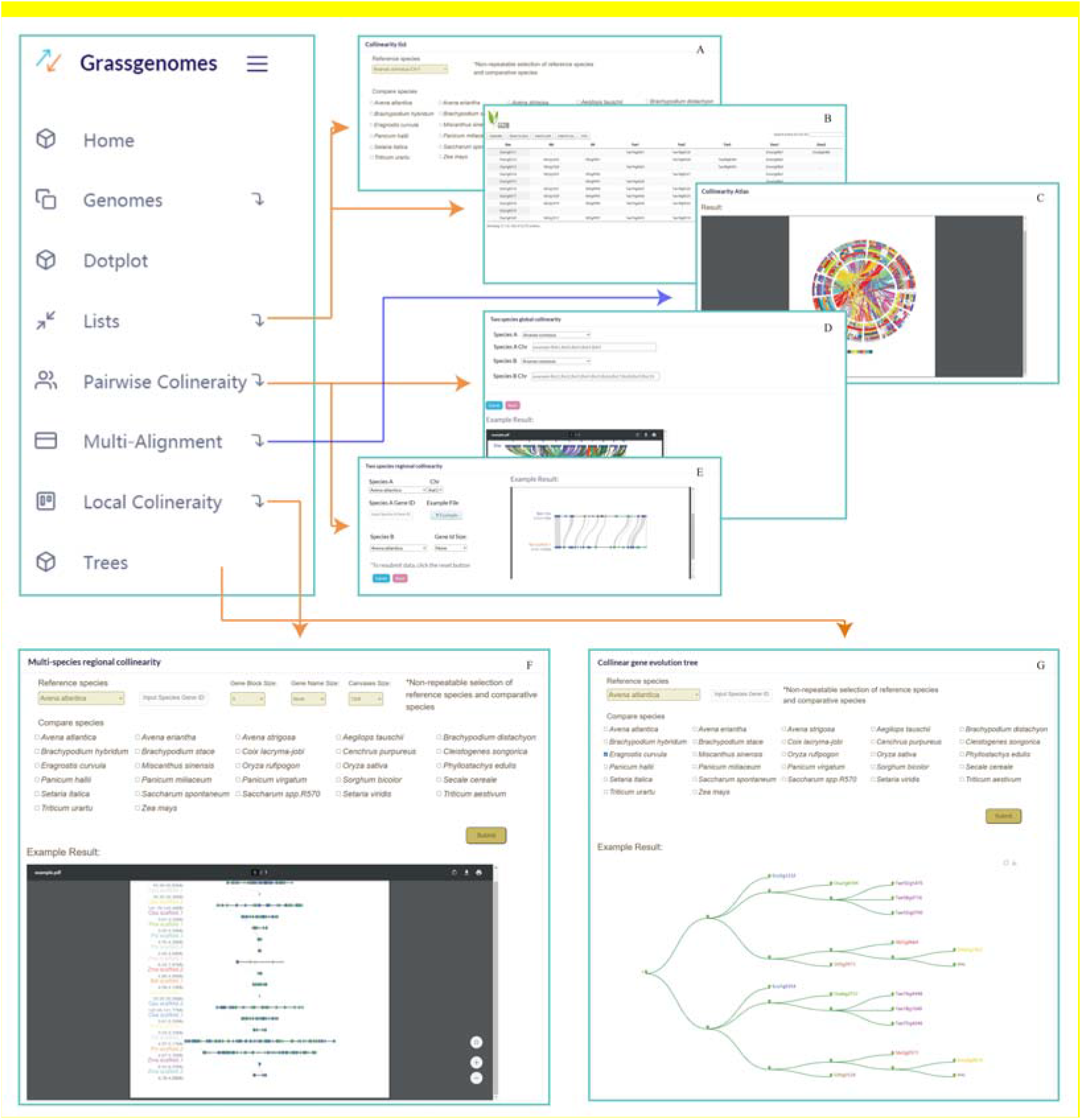
Logical relationship of modules in the database.

### Overview of data

At present, GGDB contains the information of colinear genes within a genome and between the genomes of 29 Gramineae plants. A referring outgroup, pineapple, is also included to help explore gene evolution, especially infer real evolutionary relationship of genes. As the original input data, three types of files are used: coding sequence files, protein sequence files, and general feature format (GFF) files containing chromosome sequence annotation data.

### Colinearity data

A polyploidy whole-genome duplication (WGD) event common to all main lineages of Gramineae (cWGD) was dated to 96 million years ago based on putatively neutral DNA substitution rates between duplicated genes^[53]^. Some species are further affected by another polyploid (mWGD) event. We analyzed the colinearity of pineapple and 29 species of Gramineae, made 784 comparisons between the two species, and finally generated 784 colinear lists. Then all the colinear lists are summarized into colinear tables with reference to 29 species.

Based on homologous correspondence and colinearity analysis between genomes, we obtained the homologous information of various species of Gramineae (Table 2; Supplemental table 1-4). The homologous regions are detected under the threshold of 5, 10, 20, and 50 colinear genes, respectively. Paralogous and orthologous genes are associated with polyploidy events. For example, There are 20763 homologous genes in maize genome, of which 13134 paralogous genes produced by the mWGD, and 7629 paralogous genes produced by the cWGD (Table 3;Supplemental table 5). Between rice and maize, there are 26947 orthologous genes, and 12600 outparalogous genes produced by the cWGD.

**Table 2.** List of orthologous information of some species of Gramineae.

|  | Bdi | Msi | Osa | Pmi | Pvi | Sbi | Sit | Svi | Tae | Tur | Zma |
| --- | --- | --- | --- | --- | --- | --- | --- | --- | --- | --- | --- |
| Bdi | \ | 30499 | 21053 | 30007 | 37465 | 19857 | 20782 | 21514 | 52530 | 13137 | 25708 |
| Msi | 2249 | \ | 27451 | 47405 | 60769 | 34149 | 29315 | 32667 | 72889 | 17040 | 41732 |
| Osa | 1144 | 2133 | \ | 32177 | 38906 | 21147 | 21397 | 22103 | 49571 | 12159 | 26025 |
| Pmi | 1397 | 3279 | 1236 | \ | 56986 | 31625 | 31097 | 29265 | 63541 | 16961 | 36356 |
| Pvi | 2654 | 5401 | 2547 | 3910 | \ | 42055 | 43185 | 45356 | 92787 | 23668 | 54055 |
| Sbi | 1327 | 2132 | 1068 | 1238 | 2456 | \ | 17520 | 21621 | 47707 | 11872 | 29369 |
| Sit | 1434 | 2269 | 1184 | 1345 | 2516 | 1353 | \ | 8954 | 47686 | 11494 | 27314 |
| Svi | 1481 | 2358 | 1227 | 1425 | 2764 | 1440 | 1398 | \ | 48844 | 12102 | 28198 |
| Tae | 2548 | 5482 | 2470 | 3560 | 6457 | 2311 | 2515 | 2630 | \ | 54123 | 40335 |
| Tur | 1406 | 2248 | 1345 | 1810 | 2991 | 1380 | 1342 | 1432 | 2979 | \ | 17563 |
| Zma | 1662 | 3265 | 1465 | 1960 | 3931 | 1590 | 1574 | 1762 | 3591 | 1822 | \ |

Note: Above the diagonal is the number of orthologs genes, and below the diagonal is the number of orthologs blocks.

**Table 3.**
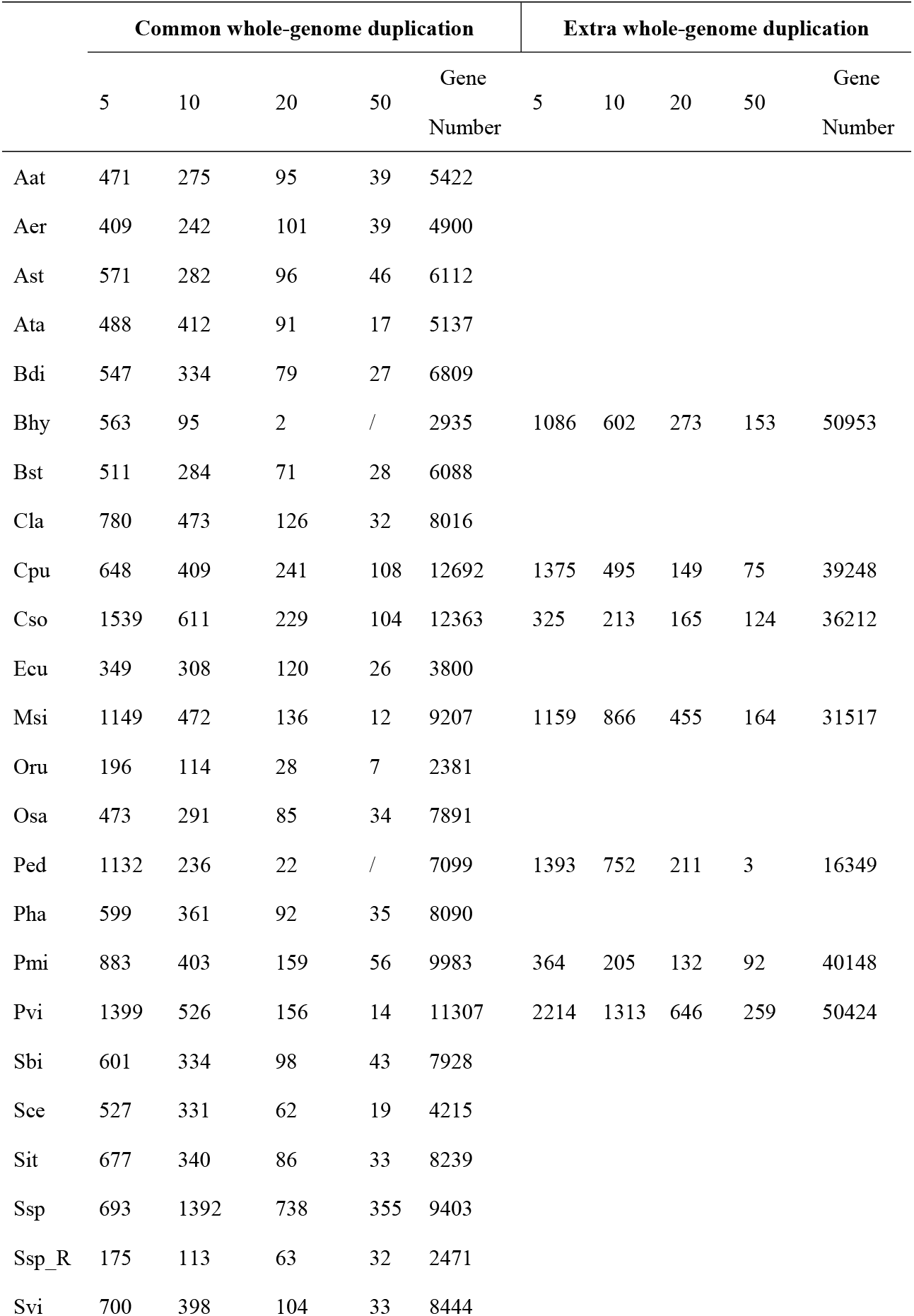
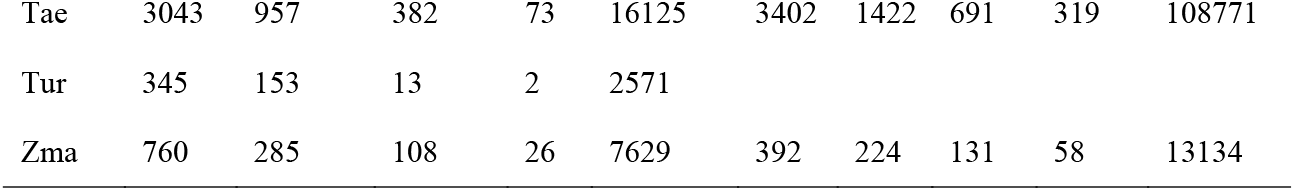
Homologous blocks produced by different whole-genomc duplication events.

|  | events |  |  |  |  |  |  |  |  |  |
| --- | --- | --- | --- | --- | --- | --- | --- | --- | --- | --- |
|  | Common whole-genome duplication |  |  |  |  | Extra whole-genome duplication |  |  |  |  |
|  | 5 | 10 | 20 | 50 | Gene<br>Number | 5 | 10 | 20 | 50 | Gene<br>Number |
| Aat | 471 | 275 | 95 | 39 | 5422 |  |  |  |  |  |
| Aer | 409 | 242 | 101 | 39 | 4900 |  |  |  |  |  |
| Ast | 571 | 282 | 96 | 46 | 6112 |  |  |  |  |  |
| Ata | 488 | 412 | 91 | 17 | 5137 |  |  |  |  |  |
| Bdi | 547 | 334 | 79 | 27 | 6809 |  |  |  |  |  |
| Bhy | 563 | 95 | 2 | / | 2935 | 1086 | 602 | 273 | 153 | 50953 |
| Bst | 511 | 284 | 71 | 28 | 6088 |  |  |  |  |  |
| Cla | 780 | 473 | 126 | 32 | 8016 |  |  |  |  |  |
| Cpu | 648 | 409 | 241 | 108 | 12692 | 1375 | 495 | 149 | 75 | 39248 |
| Cso | 1539 | 611 | 229 | 104 | 12363 | 325 | 213 | 165 | 124 | 36212 |
| Ecu | 349 | 308 | 120 | 26 | 3800 |  |  |  |  |  |
| Msi | 1149 | 472 | 136 | 12 | 9207 | 1159 | 866 | 455 | 164 | 31517 |
| Oru | 196 | 114 | 28 | 7 | 2381 |  |  |  |  |  |
| Osa | 473 | 291 | 85 | 34 | 7891 |  |  |  |  |  |
| Ped | 1132 | 236 | 22 | / | 7099 | 1393 | 752 | 211 | 3 | 16349 |
| Pha | 599 | 361 | 92 | 35 | 8090 |  |  |  |  |  |
| Pmi | 883 | 403 | 159 | 56 | 9983 | 364 | 205 | 132 | 92 | 40148 |
| Pvi | 1399 | 526 | 156 | 14 | 11307 | 2214 | 1313 | 646 | 259 | 50424 |
| Sbi | 601 | 334 | 98 | 43 | 7928 |  |  |  |  |  |
| Sce | 527 | 331 | 62 | 19 | 4215 |  |  |  |  |  |
| Sit | 677 | 340 | 86 | 33 | 8239 |  |  |  |  |  |
| Ssp | 693 | 1392 | 738 | 355 | 9403 |  |  |  |  |  |
| Ssp_R | 175 | 113 | 63 | 32 | 2471 |  |  |  |  |  |
| Svi | 700 | 398 | 104 | 33 | 8444 |  |  |  |  |  |
| Tae | 3043 | 957 | 382 | 73 | 16125 | 3402 | 1422 | 691 | 319 | 108771 |
| Tur | 345 | 153 | 13 | 2 | 2571 |  |  |  |  |  |
| Zma | 760 | 285 | 108 | 26 | 7629 | 392 | 224 | 131 | 58 | 13134 |

### Overview of the interface

On the home page of GGDB, we marked out the geographical originating locations of 29 Gramineae species on a world map ^[54]^. An interactive evolution tree including the above species is also provided on the home page. The chart interface on the home page provides interactive view of chromosomes from all species, including the numbers and lengths of chromosomes from each species and the numbers of genes on each chromosome. In addition, we use bar charts and line charts to display the chromosome numbers of each species, which makes it easier for users to compare their differences. These interactive charts can be downloaded.

### Species information page

We provide a web page for each Gramineae species, showing basic information about its name (Latin name, common name, Chinese name), picture, classification, profile (geographical distribution,biological characteristics, living habits), genome information (genome size, chromosome information, number of genes, numbers of genes located on chromosomes, number of scaffolds, length of scaffold N50), etc. We provide hyperlinks to sequencing literature for each species. Users are allowed to download DNA sequence files, protein sequence files, and general feature format (GFF) files. These species web pages can shorten the data collection time for researchers, and the format-consistent files provides convenience for following genomics research.

### Homologous gene dotplotting

Homologous gene dotplotting module is provided to show chromosome-level homology within a genome or between genomes. A homologous dotplot can be directly derived from the BLAST result (Fig. 3A), which contains relatively full information of genomic homology, as compared to the other dotplots shown below. A color scheme for gene-pair dots is adopted to separate the best-matched, often representing orthologs while comparing different genomes, and secondarily matched, and the other matched homologs.

**Figure 3.**
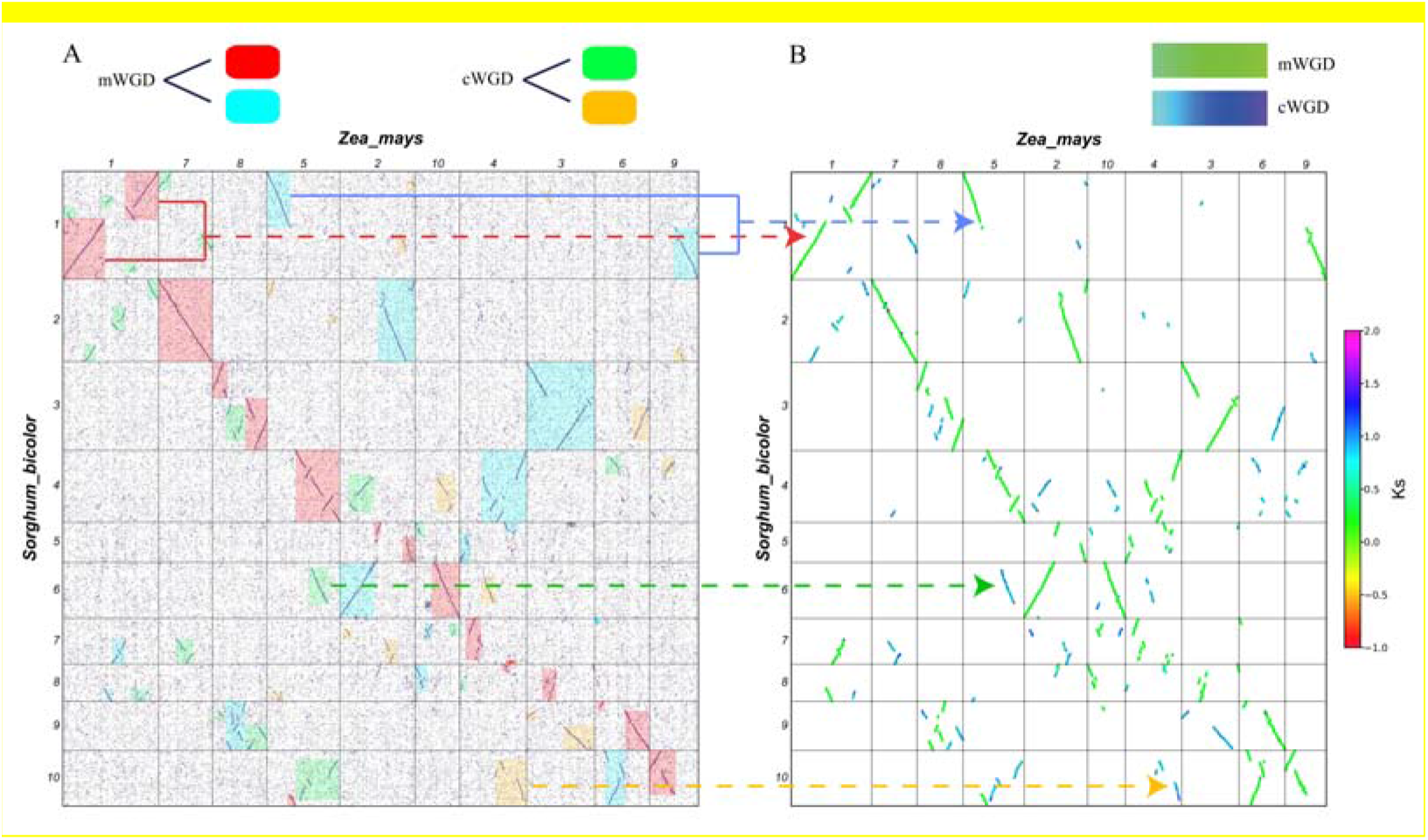
Evolutionary Analysis of homologous correspondence between sorghum and maize genomes. A. Homologous gene dotplot with blocks related to divergent ancestral whole-genome duplication events, the Grameneae-common one (cWGD) and the maize-specific one (mWGD); B. Inferred gene colinearty blocks, colored as the synonymous nucleotide substitution rates (Ks) and related to divergent whole-genome duplication events by colored arrows.

A homologous gene dotplot can be drawn by using information of inferred colinear genes and the Ks values between them (Fig. 3B). A color-scheme representing varied Ks values makes it easy to separate colinear blocks produced by different evolutionary event. Owing to the mWGD, a sorghum chromosome corresponds to two overlapping maize chromosome regions. In the meantime, it matched the other rather smaller homologous regions in maize produced due to the cWGD. Through the comparison of multiple sets of data, it is found that the cWGD of Gramineae can be distinguished from the extra whole-genome duplication in Ks=0.65. According to the Ks value of the regions, the four homologous regions of maize were divided into two groups corresponding to different whole-genome duplication events, and each group of regions was divided in detail according to the continuity of chromosome regions.

### Ks distribution

We provide a module to show Ks distribution between colinear genes. The peak of Ks between colinear genes in each species can help identify WGD event(s), for example, two Ks peaks produced be the maize colinear genes correspond to the occurrence of two WGD events affecting its evolution. The Ks peaks of colinear genes between genomes correspond to the occurrence time of species differentiation and more ancient WGD event(s), e.g., the two peaks of Ks in sorghum-maize colinear genes correspond to their speciation event and the cWGD, respectively.

### Tools

This section includes tools for comparative genomics analysis to visualize gene colinearity and phylogeny.

### Pairwise gene colinearity

The pairwise gene colinearity module can show gene colinearity at the chromosome level or at the gene level. In the chromosome level module, users select reference species and the other compared species, to find their chromosome-level homology. The colinearity at the chromosome level can help infer whether the species has experienced WGDs and/or distinguish the duplicated genes produced by events. In the gene level module, users submit an interested gene ID, select the reference species, and the other compared species, to produce an alignment of local regions. This can help find genomic changes, homologous/neighboring gene loss, DNA inversion, etc, related to the interested gene.

### Multi-genome colinearity list

The multi-genome colinear list module can produce the colinear lists generated for the species under study. There are two types of lists, with one type showing only orthologous genes between species and the other also including (out)paralogs. Users can select the reference genome and the other genomes to map onto the former to show multiple-genome alignment. In addition, we provide a variety of export formats, including excel, pdf, and csv, and copy and print functions.

### Multi-genome colinearity alignment map

The multi-genome colinearity map module is used to produce the alignment map for selected species. Users can choose any of the 29 species as reference species for comparative analysis, and maps can also be in two types, corresponding to colinearity lists shown above. With the module, a joint multi-species genome alignment map can be drawn (Fig. 4B).

**Figure 4.**
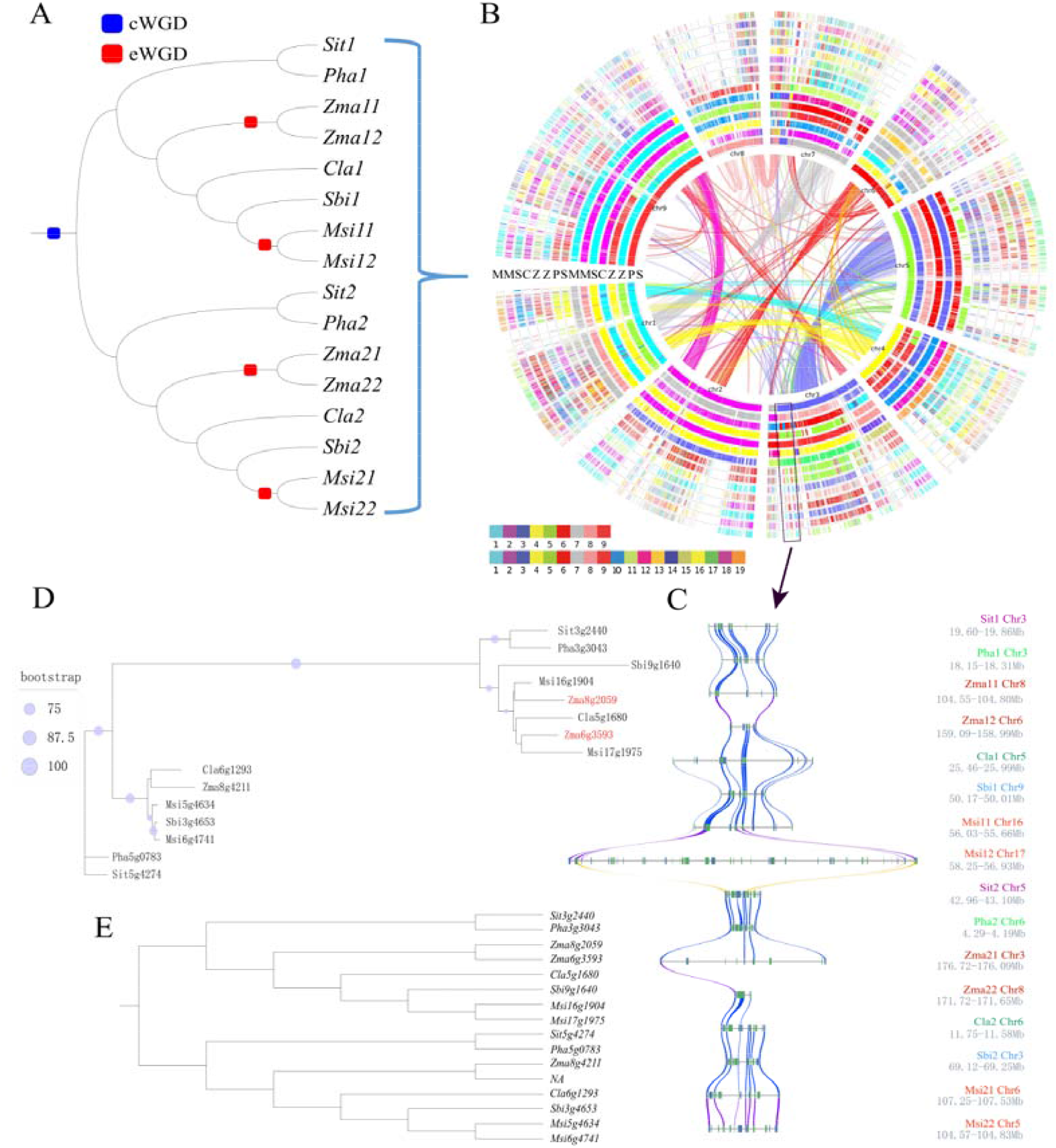
Multi-genome colinearity analysis. A. An expected gene tree of colinear homologs from selected species. Besides a common whole-genome duplication (cWGD), some species have been further affected by extra whole-genome duplications (eWGDs); *Sateria italica* (Sit), *Panicum hallii* (Pha), *Zea mays* (Zma), *Sorghum bicolor* (Sbi), *Miscanthus sinensis* (Msi), and *Coix lachryma* (Cla) are involved; B. Multi-genome alignment at a genome level; C. Multi-genome alignment in local homolgous regions with millet as the reference; D. A tree based on pure sequence alignment; E. A corrected tree based on by gene colinearity information.

### Multi-genome local colinearity

The multi-genome local colinearity module provides the function of drawing colinear maps in homologous regions of multiple species (Fig. 4C). Users select a reference species, enter an interested gene ID in the colinear list of the reference species, select the species names that needs to be compared to the reference species, and then generate a PDF format figure.

### Homologous gene evolution tree

The homologous gene evolution tree module can construct a gene evolutionary tree corrected by gene coinearity information. The user selects the reference species and a gene ID, selects the species to compare to. Two gene trees will be generated at the resulting interface, a tree based on pure sequence alignment, and the other one corrected by gene colinearity(Fig. 4D; Fig. 4E).

Actually, we found 46% of maize genes have elevated their evolutionary rates and resulted in weird phylogeny. By retrieving maize genes and their orthologous and colinear genes from sorghum, foxtail millet, rice, and weeping lovegrass (taken as outgroup), we constructed 7014 evolutionary trees and found that in 46% (3231) trees the maize genes seemed to have elevated rates (Supplemental fig. 1). In trees with one mWGD paralog, with the other one likely lost or relocated to other genomic regions, 38% (1778) showed elevated rates. In 1453 trees with two mWGD paralogs, 29% have both genes to have elevated rates and 69% to have only one gene to have elevated rates. These findings may be explained by the instability of the maize genome after the mWGD.

### Help interface

In the help interface, we provide the researcher with a detailed GGDB user manual. Users can view detailed parameter descriptions and instructions for each function in this interface.

## DISCUSSION

Grameneae plants have been recursively affected by polyploidization, which makes their genome much complex to decipher^[55; 56]^. Owing to this fact, gene collinearity inference in present databases is not well related to each polyploidization event, which holds back the research to understand gene function origination and innovation. Actually, the appearance and establishment of novel gene functions can often be related the production and divergence of duplicated genes, the most of which were produced by ancestral polyploidization^[57; 58]^. Here, by separating duplicated genes produced by different polyploidization, and separating orthologous genes from outparalogous when comparing different species, we set up the GGDB to store event-related collinear genes in 29 Grameneae plants and one outgroup. The related work is intense in that 420 pairwise comparisons have been done between genomes and eventually hierarchically constructed multiple comparisons by selecting reference genomes in each major groups, subfamilies or genus. To separate homologs produced by different events involved, computational analysis of their divergence and artificial identification of complement homologous blocks aroused by chromosome breakages were performed.

The present database provided friendly tools for the users to show pairwise or multiple genome alignment in the global or local levels, and produce lists of homologous genes, related to evolutionary events, which provides opportunities to perform deep study of their evolution and functional innovation. Figures of homologous gene dotplots, alignment of selected genomes, and evolutionary trees of selected genes can be downloaded for further research on genome structural changes, chromosome rearrangements, gene losses, and gene functional evolution.

In that different copies of homologous genes have divergent evolutionary rates^[59; 60]^, we provided a module to correct evolutionary trees constructed purely using sequence alignment, by using information of gene colinearity and shared evolutionary events. Actually, elevated evolutionary rates were observed in considerable percentages of paralogous genes produced by polyploidization^[48; 53]^, which often resulted in aberrant evolutionary trees that cannot be corrected by selecting methods to construct the trees. The tree correction module provides a means to make a realistic tree for the researchers to manipulate for their further study of gene evolution and functional innovation.

In the future, with more and more genome sequences released, we will continue to add new genome data in the GGDB. We also encourage users to submit their new Gramineae sequencing data sets to the GGDB to enrich and improve the database. GGDB will act as a comparative genomics platform for genomics research of Gramineae and the other related Monocotyledons.

## AVAILABILITY

**The GGDB can be accessed through the web server at http://www.grassgenome.com/**.

